# Multi-Antigenic Virus-like Particle of SARS CoV-2 produced in *Saccharomyces cerevisiae* as a vaccine candidate

**DOI:** 10.1101/2020.05.18.099234

**Authors:** Kajal Arora, Ruchir Rastogi, Nupur Mehrotra Arora, Deepak Parashar, Jeny Paliwal, Aelia Naqvi, Apoorva Srivastava, Sudhir Kumar Singh, Sriganesh Kalyanaraman, Swaroop Potdar, Devanand Kumar, Vidya Bhushan Arya, Sarthi Bansal, Satabdi Rautray, Indrajeet Singh, Pankaj Surendra Fengade, Bibekanand Kumar, Prabuddha Kumar Kundu

**Affiliations:** Premas Biotech Private Limited, Plot 77, Sector 4, IMT Manesar, Gurgaon 122050, Haryana, India

## Abstract

Spike, Envelope and Membrane proteins from the SARS CoV-2 virus surface coat are important vaccine targets. We hereby report recombinant co-expression of the three proteins (Spike, Envelope and Membrane) in a engineered *Saccharomyces cerevisiae* platform (D-Crypt™) and their self-assembly as Virus-like particle (VLP). This design as a multi-antigenic VLP for SARS CoV-2 has the potential to be a scalable vaccine candidate. The VLP is confirmed by transmission electron microscopy (TEM) images of the SARS CoV-2, along with supportive HPLC, Dynamic Light Scattering (DLS) and allied analytical data. The images clearly outline the presence of a “Corona” like morphology, and uniform size distribution.

## Introduction

SARS-CoV-2,, has become a global pandemic in last five months and is posing a grave challenge on many fronts, including health, economy, travel and social norms of living(*1*). The incidence of mortality and morbidity observed with this pandemic, along with the widespread of disease has created an immediate and urgent need for focused steps to develop a vaccine as an earliest measure that can be utilized for masses (*2*). Many candidates are being developed for testing as a vaccine for SARS-CoV-2 and research data along with prior knowledge of technology behind each candidate becomes key resource to understand the mechanism of demonstrating immunogenicity and efficacy. Amongst different technologies that are being adapted for developing the vaccine candidate, VLPs are known to be a potent technology for increased immunogenicity and there are number of examples of commercial vaccines and candidates under trial as VLPs (*3, 4*). Most of the VLP vaccines utilize a major surface protein to form the VLP (*5*). A strong B cell response is shown by VLPs even in the absence of adjuvants due to efficient crosslinking with specific receptors on B cells (*6, 7*). In developing VLPs as vaccine candidates, one of the significant challenges has been to incorporate multiple surface antigens, express them in a single host and allow self-assembly into a VLP (*5*).

In this study, we report the first successful self-assembly of the SARS CoV-2 VLP by co-expression of the S, E and M proteins in *Saccharomyces cerevisiae* expression platform. The VLPs were studied by multiple analytical methods, and imaged using transmission electron microscopy (TEM). The TEM images confirmed the morphology and their self-assembly into a VLP.

The coronavirus family is known to comprise of four major structural proteins in its genome, namely the spike (S) protein, membrane (M) protein, envelope (E) protein and the nucleocapsid (N) protein. All of these proteins are encoded within the 3’ end of the genome (*8*). The S protein is known to mediate attachment of the virus to the host cell surface receptors resulting in fusion and subsequent viral entry into the host cell (*8*). The M protein is the most abundant protein and defines the shape of the viral envelope (*8*). The E protein is the smallest protein of all the structural proteins in the viral capsid and its function is described in viral assembly and budding (*8*). The N protein binds to the RNA genome and also functions along with M protein in viral assembly and budding. As a class of subunit vaccines, the virus-like particles (VLPs) are differentiated from soluble recombinant antigens by stronger protective immunogenicity, associated with their structure, and hence make a much more relevant vaccine candidate (*9*).

The episomal constructs for the three proteins, S, E and M with C-terminus His tag were transformed into *Saccharomyces cerevisiae* expression platform independently and analyzed for expression in the 24^th^ h post induction. His tag was used to enable detection of the expressed proteins as there are no commercially available antibodies against the proteins S, E and M. Immunoblot analysis confirmed the expression of the three proteins using anti-His antibody with specific band at ~26 kDa and ~9.5kDa for M and E protein respectively (Figure 1) which was also confirmed by mass spectrometric analysis. However, a band at ~80kDa was observed for S protein (Figure 1) which is due to detection of the S2 domain of S protein. The S protein is known to be cleaved by a host cell protease into polypeptides S1 and S2 in yeast (*10*). While the band for the cleaved S2 protein fragment is expected at ~66kDa, we obtained a higher molecular weight band, which may be possibly due to the fact that the protein is known to be glycosylated. The expected molecular weight of the S protein is ~ 140 kDa. Interestingly, peptide mapping data showed presence of the complete S protein sequence, matching to full length 140 kDa protein, suggesting cleavage of S protein by the cellular protease into its two fragments, S1 and the S2, which were found to be coexisting together on the microsomal membrane, and would would get incorporated into the a VLP as it assembles at the ER-Golgi interface.

**Figure 1:**
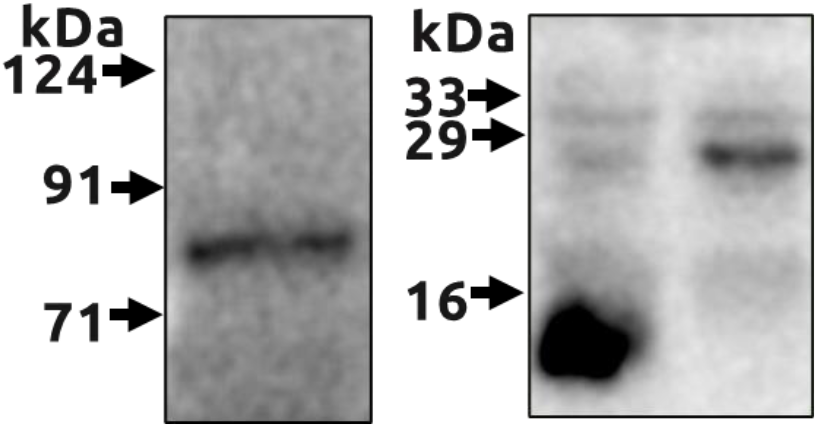
Confirmation of expression of Spike, Envelope and Membrane protein in S. cerevisiae

**Figure 2:**
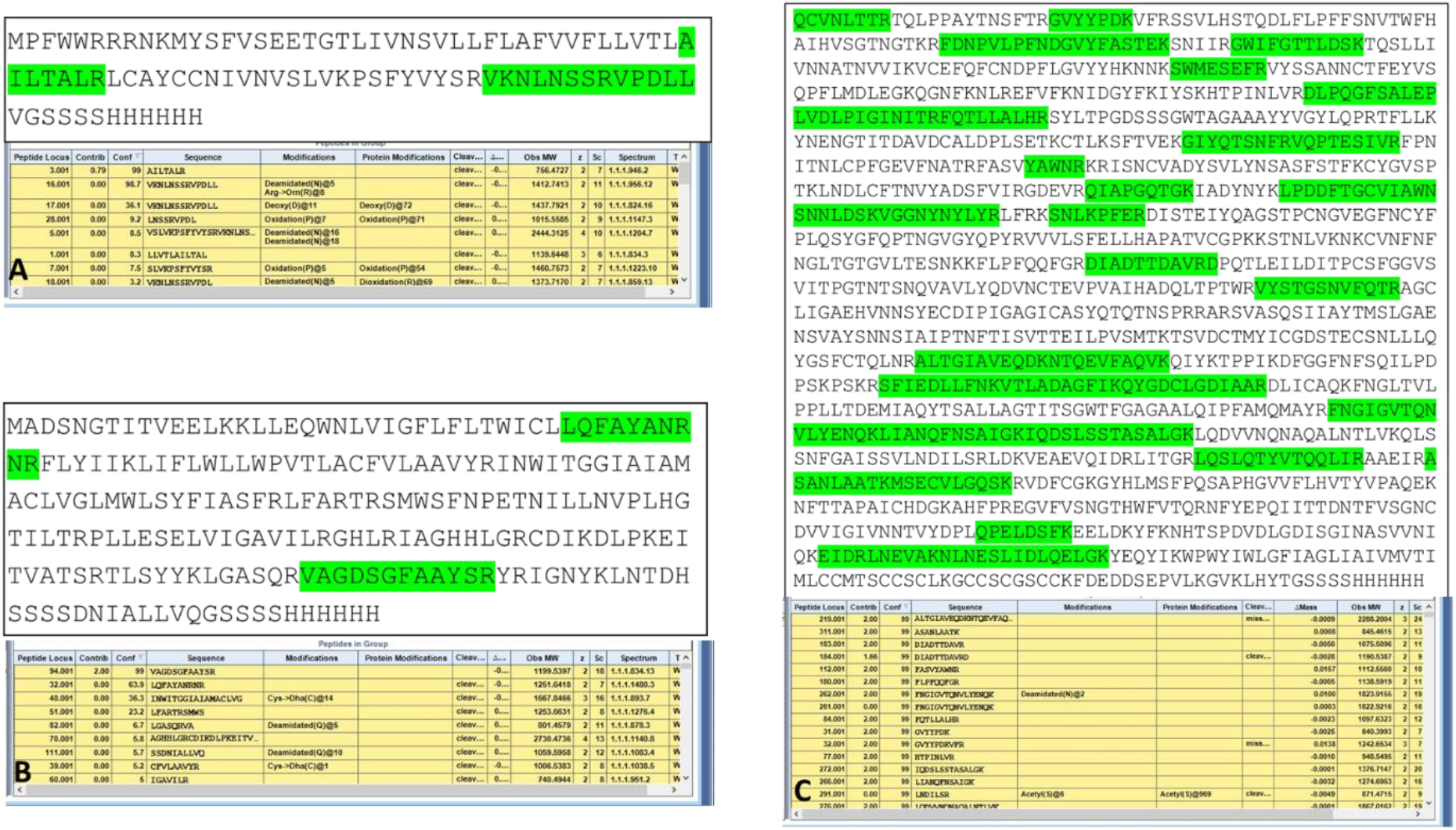
Peptide mapping data confirming the expression of Envelope (A), Membrane (B) and the Spike (C) proteins

In order to co-express the S, E and M proteins into *S. cerevisiae*, recombinant construct for the expression of S and E protein, pYRE100_CSP_CEP_His and M protein, pYRI100_CMP_His construct were co-transformed into the host and the positive clones were selected on YNB-URA-LEU plates. Selected clones were taken forward for expression analysis. Expression of the three proteins, S, E, and M were detected using anti-His antibody under both reducing and non-reducing conditions. Results in Figure 3 show the expression of S, E and M proteins. We do observe differential expression of the three proteins. Further, a band was observed at ~33kDa which may be due to oligomerization of E and M protein. Interestingly, the band at ~71kDa disappears under non reducing conditions suggestive of formation of high molecular wt. complexes or VLPs. It has been previously demonstrated that co-expression of S, E and M has been shown to form VLPs (*10, 11*) for SARS-CoV. Importantly, the yeast expression platform along with its GRAS status provides the high scalability, robustness and cost-effective production for billions of dosages which would be the required to fight the pandemic globally.

**Figure 3:**
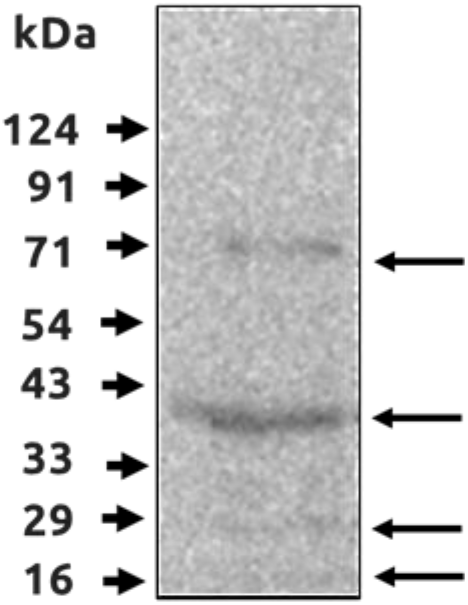
Co-expression of Spike, Envelope and Membrane protein in in S. cerevisiae (PYPD)

In order to confirm the presence and determine the morphology and size of the VLPs formed on over expression of S, E and M protein in *S. cerevisiae*, parallel HPLC (High Performance Liquid Chromatography), DLS (Dynamic Light Scattering) (*12*) and Transmission Electron Microscopy (TEM) studies were carried out. *S. cerevisiae* with vector only was analyzed as negative control. Cell lysates of various hours’ time point post induction *from S. cerevisiae yeast host*, overexpressing the three antigenic proteins were fractionated on SEC-HPLC. The high molecular weight peak was absent in the vector transformed cells (negative controls) (Figure 4A). The high molecular weight fraction was collected and subjected to analysis by DLS and TEM. The high molecular weight peak fractions were re analyzed for their purity using SEC-HPLC (Figure 4B). The samples at 72 h post induction, showed maximal concentration of the VLPs in the lysate. The samples, obtained at 72 h post induction from the lysate, were taken forward for DLS and TEM analysis. DLS analysis data showed a range of Particle size of 90 - 124 nm, with a mean of 103 ± 17 nm was observed for the VLPs which is evident from the correlation function graph (Figure 4C). This data also indicates that the VLP particles are large with poly dispersity index less than 25%. Interestingly, TEM analysis showed that VLPs were spherical in shape (Figure 5A) with distinct spike like structures decorating the outer surface (Figure 5 B, C). The outer surface was highly dense as compared to the core as observed on negative staining of the VLPs. Mass spectrometric analysis of the VLPs confirmed the presence of S. The E and M protein peptide could not be detected possibly due to their small size. However, the existence of the M protein is understood by the fact for its essentiality for formation of viral envelope (*13*).

**Figure 4.**
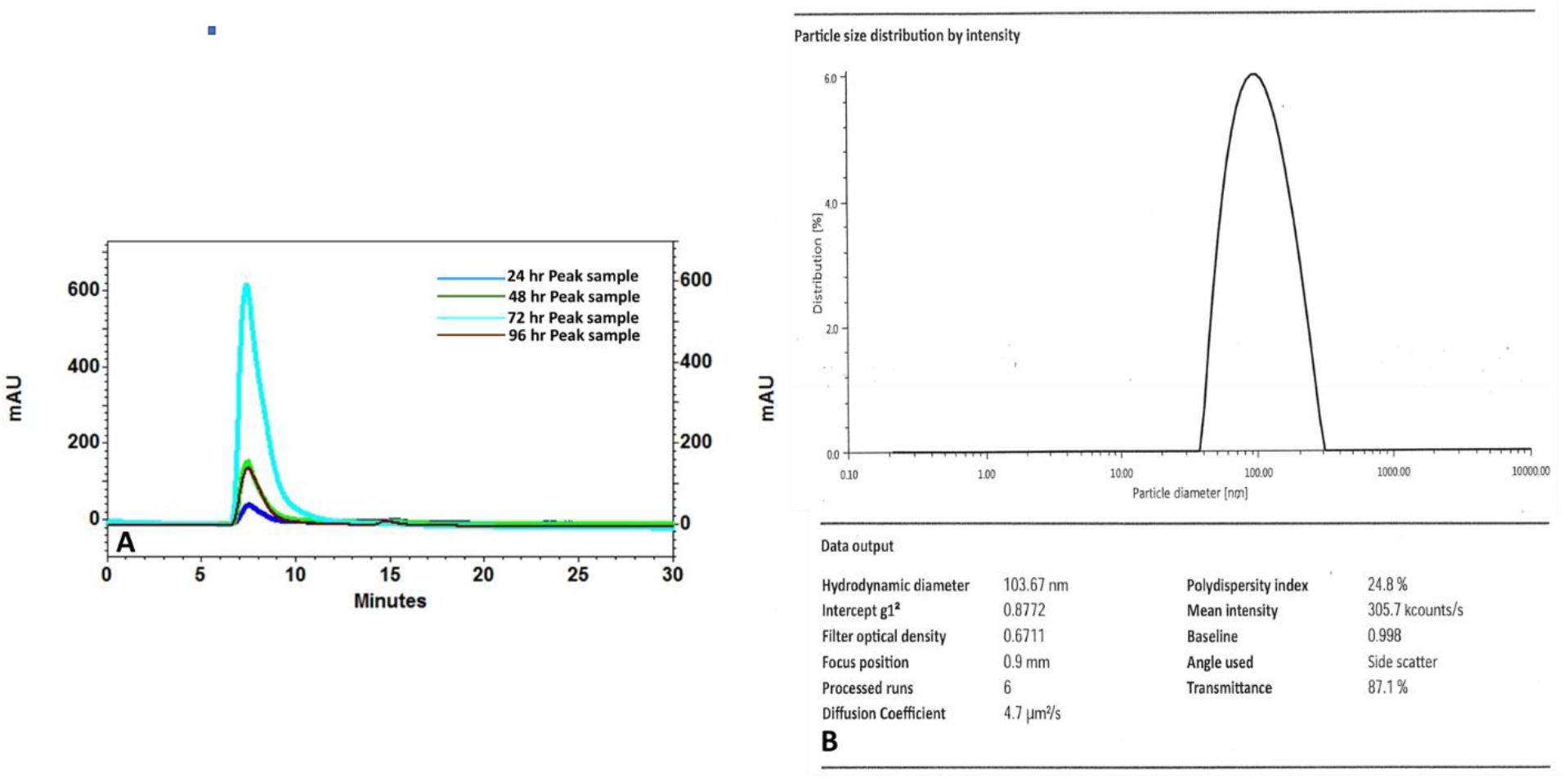
(A) Overlay of HPLC profile of purified Peak from S. cerevisiae cells cytoplasmic extract, co-expressing S, E and M protein from the time points, 24h, 48h, 72h and 96h. (B) Dynamic Light Scattering (DLS) analysis of the 72h sample

**Figure 5:**
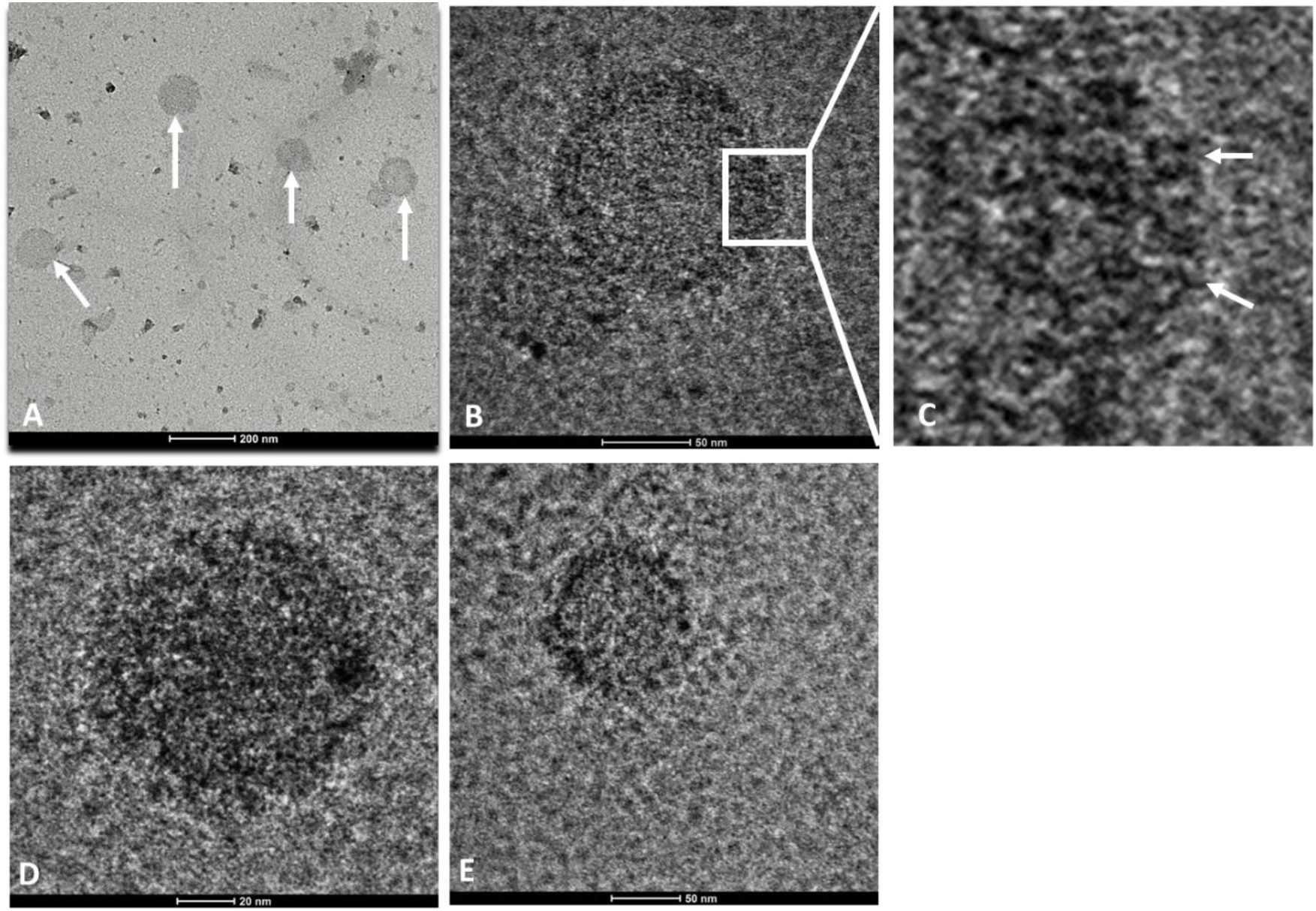
(A)TEM images of Virus Like Particle (VLP) from yeast cell cytoplasmic lysate overexpressing recombinant Spike, Envelope, Membrane proteins, 72h sample from cell lysate, (B) TEM image at 120KX magnification (C) Zoomed in image of VLP (D,E) TEM image at 400KX and 190KX magnification

This is the first time that the VLP of the SARS CoV-2 virus has been imaged under TEM and allied analytical data concurrently provide the biophysical characterization evidence of a VLP. This would enable to develop a potent vaccine candidate against SARS CoV-2 which being in yeast would be scalable, cost effective and provide a simplified production process.

## Acknowledgements

The authors thank Dr Vineeta Batra, G.P. Pant Hospital for the Transmission Electron Microscopy. We thank Dr Anurag Agarwal, Director, IGIB for his kind support and advice, Dr Archana Singh, IGIB, for her help for the TEM processing. We thank Dr Nirpendra Singh, RCB, for the Mass Spectrometry Analysis. The authors acknowledge the generous support from Premas Biotech and Akers Biosciences for co-development of this study. The work was carried through the financial support from Premas Biotech and Akers Biosciences. The authors thank Mr Gaurav Kaushik and Mr Sanjib Pathak for their regulatory support work and documentation support.

## Materials and methods

### Cloning of “S”, “E” and “M” proteins

In order to express the S, E and M proteins from SARs CoV-2 proteins were cloned into our proprietary expression vectors, pYRE100 and pYRI100 (*14*). The M protein was cloned in pYRI100 integration vector (pYRI_CMP_His) and integrated into PYPD genome while S and E (pYRE100_CSP_CEP_His) protein were cloned into pYRE100 episomal single expression vector as two separate expression cassettes. The synthetic genes sequence biased and optimised for expression in *S. cerevisiae* were obtained from GeneArt (Regensburg, Germany). Further, all the genes were synthesized so as to have a His-tag at the C-terminus of the protein for the ease of detection as no antibodies were available. All the genes were also synthesised without an His tag. As an initial proof of concept, all the three proteins with a C-terminal His tag were cloned into pYRE100 vector and transformed for expression analysis.

### Transformation in *S. cerevisiae* PYPD strain

Episomal constructs of S, E and M with His tag, pYRE100_CSP_His, pYRE100_CMP_His and pYRE100_CEP_His plasmid and host vector pYRE100 plasmid were transformed into proprietary protease deficient PYPD yeast expression strain using Lithium acetate/SS-DNA/PEG mediated protocol and transformants were selected over selective YNB Glucose minus URA plates. Also integrating construct with M protein was transformed in PYPD host and selected on YNB minus LEU plates. For co expression of three proteins, episomal construct with E and S gene were co-transformed into *S. cerevisiae* (PYPD) containing integrated M protein and transformants selected on YNB Glucose (without URA and LEU) at 28°C for 2-3 days. In order to study the expression, three isolated healthy transformed colonies were inoculated in appropriate YNB Glucose media maintaining auxotrophic selection at 28°C for 36 h. A colony of host strain PYPD transformed with pYRE100 served as host-vector control. At this stage, cultures were harvested and the cell pellet was induced with galactose at a final concentration of 2% in YNB minimal medium without respective markers (~OD/ml at this time for all the cultures was 5). All the cultures were harvested 24^th^ h post induction and processed for Western blot analysis.

### Expression analysis

Briefly, the cells were pelleted and treated with 2M Lithium acetate on ice for 5 min. Subsequently, the cells were centrifuged at 13000 rpm for 1 min. The cell pellet was then treated with 0.4M NaOH for 5min. The cells were finally pelleted and re-suspended in reducing dye and boiled for 10 minutes. Samples corresponding to 1.4 OD600 of cells were resolved over SDS-Western blot was developed using anti-His specific antibody (Cat no. Sigma:H1029) at a dilution of 1:3000 and HRP conjugated anti-mouse secondary antibody (Anti mouse IgG-A4416) at dilution of 1:4000.

### Induction studies to enhance the biomass and VLP production

In order to increase cell biomass and the overall specific yields of the proteins, fed batch culture was carried out. The pre-seed and seed media were prepared in selective media “YNB -Ura -Leu” and finally the culture was transferred to 5L YPD media (HiVeg Peptone 20gm/l; Yeast extract 10gm/l and Dextrose 20gm/l). The key fermentation parameters of temperature (28 °C), pH (5.8), D.O (30%) were maintained automatically. The culture was induced at 24 hr with 125 ml of 40% Galactose along with 10X YT (Yeast extract 5gm/L and Tryptone 10gm/L). The culture was fed with 40% galactose and 10X YT every 24 h.

### SEC-HPLC analysis cell lysates

In order to study the expression of the overexpressed S, E and M protein, Immuno-blot blot analysis was carried out using the cell lysate prepared after disruption of cells using glass beads.

SEC HPLC fractionation: 10 μL of both cell lysate samples at different hours of fermentation were injected onto TSKgel SuperSW 2000 (4.6 mm ID × 30 cm L (Tosoh Bioscience GmbH)) column for analytical SEC analysis and analysed with Agilent Technologies HPLC instrument (1260 Infinity II). The estimated exclusion limit of this column for proteins is 150,000 Da for globular proteins. 20 mM sodium phosphate, pH 7.0 with 400 mM sodium chloride was used as a standard SEC buffer for the elution which provides necessary ionic strength to the VLPs. UV signals were traced at 214 nm and 280 nm. 0.25 mL/min was the flow rate used for the analysis. Corresponding VLP peak was separated by injecting 100 μL into the chromatograph and peaks were collected from the detector end and further used for DLS or TEM analysis.

### Dynamic Light Scattering Analysis

Dynamic light scattering analysis was done as a first proof of concept to see the VLP formation from the shake flask culture using Litesizer 500 (Anton Paar, Austria) instrument. The size of the molecule was measured as the function of maximum intensity observed. The laser used to measure the hydrodynamic diameter had the wavelength of 658nm. During the measurement, for the light scattering angle, automatic mode was selected which automatically selects the angle based on the particle concentration. In this case, side scattering (90°) was used by the instrument to measure the Hydrodynamic diameter.

### Electron microscopy

In order to characterise and determine the size and structure of VLPs, the HPLC purified fractions from the cell ly sate were subjected to electron microscopic analysis as described previously (*15*). Briefly, the fractions were adsorbed onto Carbon-Formvar copper grids and subsequently negatively stained with 1.5% uranyl acetate aqueous solution for 50s. Subsequently the grids were washed and examined on Talos Arctica transmission electron microscope.

## Notes

### Competing Interest Statement

The authors have declared no competing interest.

